# Reversion to metabolic autonomy underpins evolutionary rescue of a bacterial obligate mutualism

**DOI:** 10.1101/2024.06.27.600993

**Authors:** Ignacio J. Melero-Jiménez, Yael Sorokin, Ami Merlin, Alejandro Couce, Jonathan Friedman

**Author notes:** Corresponding author. (I.J.M.J); (A.C.); (J.F.). These authors contributed equally to this work (Alejandro Couce, Jonathan Friedman).

## Abstract

Populations facing lethal environmental change can avoid extinction by undergoing rapid genetic adaptation, a phenomenon termed evolutionary rescue. While this phenomenon has been the focus of much theoretical and empirical research, our understanding of evolutionary rescue in communities consisting of interacting species is still limited, especially in mutualistic communities, where evolutionary rescue is expected to be constrained by the less adaptable partner. Here, we explored empirically the likelihood, population dynamics, and genetic mechanisms underpinning evolutionary rescue in an obligate mutualism in which auxotrophic *Escherichia coli* strains exchanged essential amino acids reciprocally. We observed that >80% of the communities avoided extinction when exposed to two different types of lethal and abrupt stresses. Of note, only one of the strains survived in all cases. Genetic and phenotypic analyses show that this strain reverted to autonomy by metabolically bypassing the auxotrophy, but we found little evidence of specific adaptation to the stressors. Crucially, we found that the mutualistic partners were substantially more sensitive to both stresses than prototrophs, so that reversion to autonomy was sufficient to alleviate stress below lethal levels. We observed that increased sensitivity was common across several other stresses, suggesting that this may be a general property of obligate mutualisms mediated by amino acid exchange. Our results reveal that evolutionary rescue may depend critically on the specific genetic and physiological details of the interacting partners, adding rich layers of complexity to the endeavor of predicting the fate of microbial communities facing intense environmental deterioration.

## Main

Environmental changes can lead to species becoming maladapted and decline toward extinction, but sometimes extinction can be avoided if species adapt rapidly, a phenomenon termed evolutionary rescue^1^. Much effort has been dedicated to elucidating the factors that affect the likelihood and dynamics of evolutionary rescue as it is central to several important areas including conservation, agriculture and medicine. For example, in conservation, evolutionary rescue may inform our efforts to maintain biodiversity in the face of phenomena like habitat loss or climate change^2,3^. In contrast, in the fields of medicine and agriculture, evolutionary rescue can thwart our efforts to eliminate drug-resistant pathogens and pests^1,4,5^.

Previous studies have focused on several major factors that determine the likelihood of evolutionary rescue, including the rate of environmental change (abrupt vs. gradual), dispersal rate or population size^1,4,6–9^. However, all of these studies have focused on populations composed of a single strain, whereas natural populations typically harbor multiple interacting strains and species. Interestingly, these interactions are predicted to alter the likelihood of evolutionary rescue^10^. Indeed, theory suggests that both competition^11,12^ and cooperation^13,14^ can have significant impact on evolutionary rescue dynamics. These effects are mediated by both demographic and genetic effects, as interactions can alter population sizes and introduce additional selective pressures and adaptive constraints^15,16^. Moreover, interactions can also affect adaptation to changing conditions by altering diverse population genetics parameters experienced by the interacting partners, as recently observed with a toxin from *Burkholderia cenocepacia*, which has mutagenic effects in the competing species it fails to kill^17^.

An important knowledge gap is how evolutionary rescue is affected by mutualistic interactions, which play a key role in shaping communities in nature. Theoretically, evolutionary rescue of obligate mutualisms is expected to be challenging since the survival of the system is determined by the least adapted species. In this light, the weakest link hypothesis^18,19^ posits that the adaptation rate is slower in mutualisms than in isolation, because while a single strain can become adapted with a single mutation, mutualistic populations require multiple mutational events to achieve the adaptation of all partners. Besides, stress can lead to a weakening of the mutualistic interactions^18^, eventually leading to mutualism breakdown^20^. Examples of mutualism breakdown have been observed in natural settings, as in the interaction between corals and/or anemones and *Symbiodiniaceae* (i.e., bleaching)^21^ and in the human-induced disturbance of soil microbiome^22^. Under more controlled conditions, a recent study demonstrated that obligate mutualistic *Escherichia coli* communities have a lower likelihood of survival to increasing levels of antibiotics and that they often break down and revert to autonomy^23^.

Intrigued by these results, we aimed to empirically gain further insights into how obligate mutualisms can avoid extinction when exposed to abrupt stresses, which are considered to pose the most challenging scenario for evolutionary rescue in general^7,24^. As predicted by the weakest link hypothesis, the challenge is exacerbated in obligate mutualisms, since evolutionary rescue is expected to require either a rapid adaptation of both partners or one strain both adapting to the stress and reducing its dependence on the partner. Here, we exposed a synthetic two-strain *E. coli* community engaged in obligate cross-feeding to abrupt environmental change imposed by two qualitatively different stressors (Figure 1): salinity or p-nitrophenol (PNP). We chose these two treatments due to their general effect on bacterial physiology. While salinity causes hyperosmotic stress and inactivation of crucial cellular processes^25^, phenolic compounds have toxic effects on cell membranes due to their high aqueous solubility^26^. Beyond quantifying the likelihood of ER, we also sought to characterize the evolved strains both phenotypically and genotypically in order to elucidate the mechanistic underpinnings of ER in our system.

**Figure 1.**
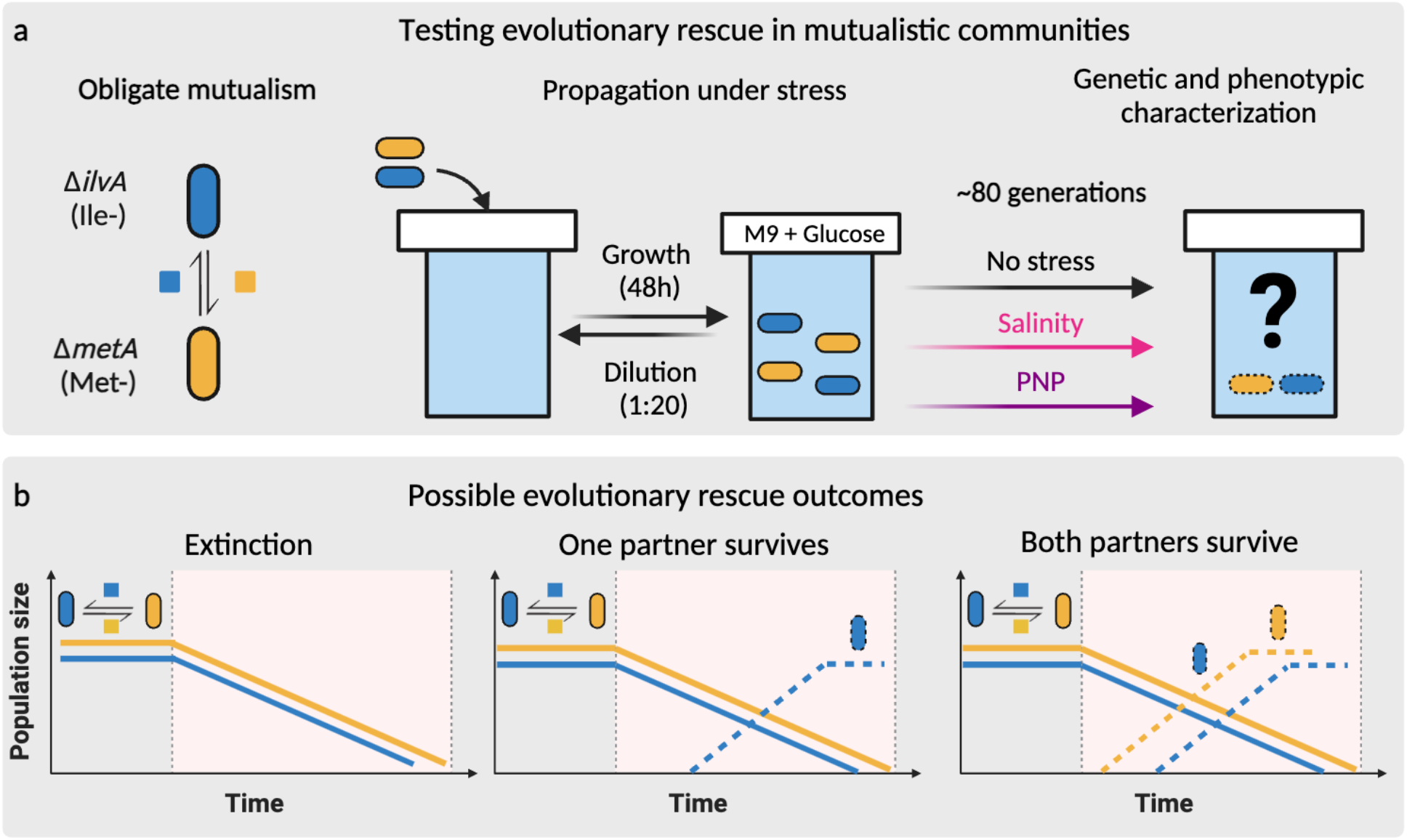
Testing how readily evolutionary rescue can occur in populations engaged in obligate mutualism. **a**, we utilized a pair of *E. coli* strains engaged in obligate mutualism based on amino acid exchange and exposed them to three treatments (No stress, salinity, and PNP) to observe if the community could avoid extinction through evolutionary rescue. **b**, Abrupt environmental stress is expected to cause a decline in population density towards extinction, followed by three possible scenarios: extinction of both partners, adaptation of one partner that recovers while the other goes extinct, or adaptation and recovery of both partners.

## Results

### Evolutionary rescue prevents the extinction of bacterial communities engaged in an obligate mutualism

To test whether obligate mutualism alters the likelihood of evolutionary rescue, we evolved communities consisting of a pair *E. coli* strains that were previously engineered to be auxotrophic to complementary amino acids^27^, as well as populations of the prototrophic *E. coli* from which these auxotrophic strains were derived (see Methods). We chose knockouts in the *ilvA* (ΔI) and the *metA* gene (ΔM), because previous work has shown that they can form a robust obligate mutualism based on methionine-isoleucine exchange (Figure S1). Next, we propagated replicate cocultures of auxotrophic strains and monocultures of the prototrophic strain for 20 growth-dilution cycles (∼80 generations) under each of three treatments (n=48 for each treatment): no stress, salinity (3%), and PNP (0.4 μM).

Without stress, both mutualistic communities and prototrophic populations maintained stable population densities (Figure 2a). In contrast, communities engaged in obligate mutualism declined towards extinction when exposed to either stress. However, despite orders or magnitude drops in population size, the majority of mutualistic communities were able to recover and survive (37 out of 48 communities survived under salinity, and 41 out of 48 communities survived under PNP), exhibiting the typical U-shaped curve associated with evolutionary rescue^4^. No significant association between the stress conditions and survival rates was observed (Chi-squared test, p-value = 0.38).

**Figure 2.**
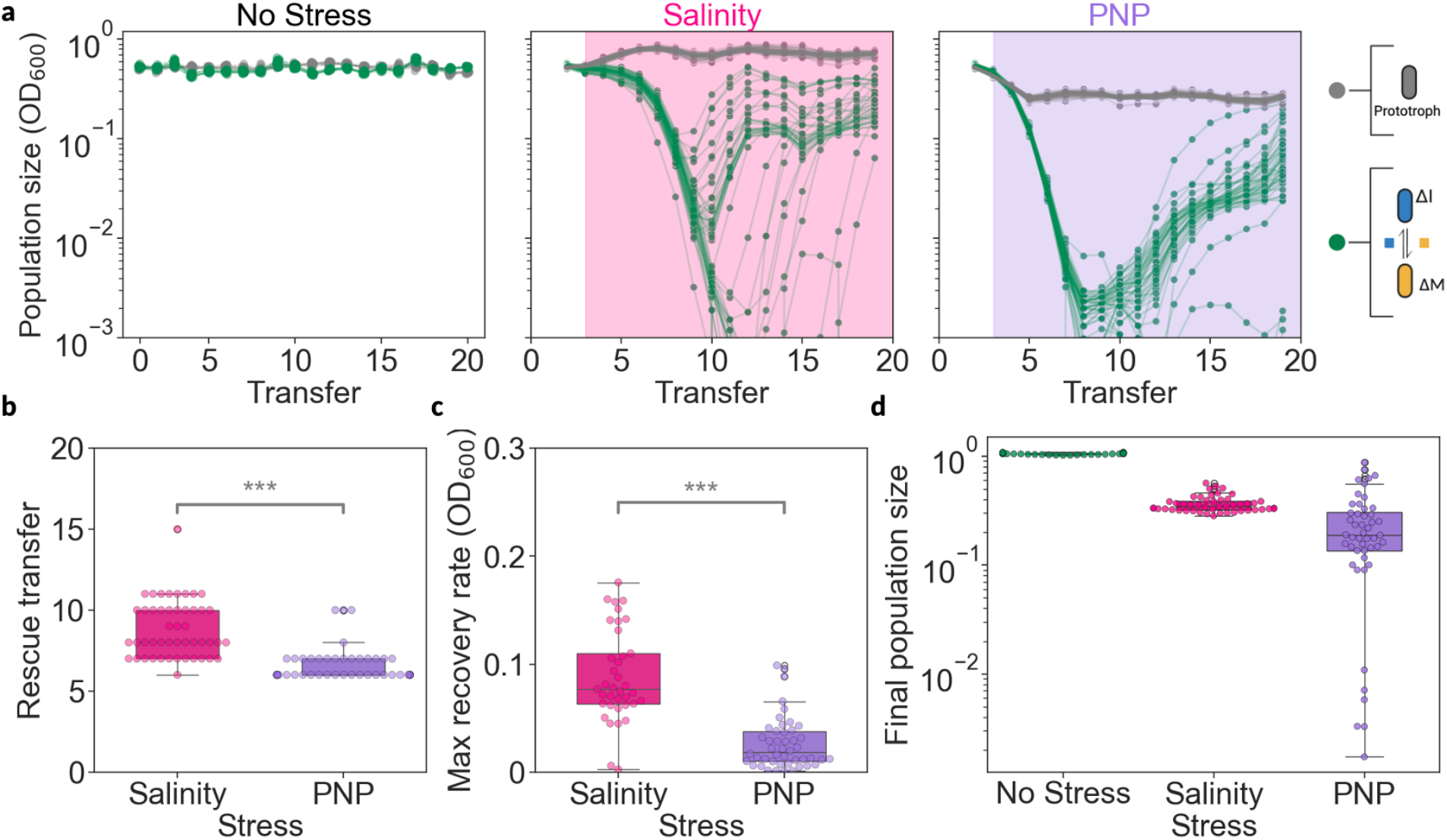
Evolutionary rescue prevents the extinction of bacterial communities engaged in an obligate mutualism based on metabolic exchange. **a**, Population dynamics of the prototrophic strain (gray) and the obligate mutualism (green) in three different stress treatments (no stress, salinity, and PNP). The red and purple background indicate exposure to salinity or PNP, correspondingly. The experiment consisted of 48 independent populations for each treatment. **b**, Box Plots representing the transfer when communities/populations begin to recover after stress exposure (OD_600_ shifts from negative to positive trend). **c**, Maximal rate of recovery computed as maximum change in population size per transfer for each experimental culture after stress exposure (OD_600_/transfer). **d**, Final population size of recovered mutualistic communities, calculated as the median of the last three transfers normalized relative to the median of the prototrophic strain at the final transfer. P values (b and c) ***<0.001 (two sided Mann–Whitney U test)

We next sought to characterize whether the different treatments and replicates display common patterns or distinct dynamics of evolutionary rescue. To this end we quantified the time it took each population to start recovering, the speed of population size recovery, and the population density they finally reached (see Methods). On average, salinity-exposed communities required 2 more transfers to recover than those exposed to PNP (Figure 2b; median of 8 *vs*. 6 transfers; two sided Mann–Whitney U test, p-value ∼ 10^−12^), but once recovered they grew significantly faster (Figure 2c; median of 0.07 *vs*. 0.03 OD_600_/Transfer; two sided Mann–Whitney U test, p-value ∼ 10^−10^) and to higher population densities, approaching those of the prototrophic strain (Figure 2d; median 0.35 *vs*. 0.19 normalized OD_600,_ two sided Mann–Whitney U test, p-value ∼ 10^−21^). Notably, the recovery dynamics were more variable in salinity-exposed communities than in PNP-exposed ones (Levene test of equality of variances, p-value ∼ 10^−43^). These findings suggest that there is a larger variability in the genetic changes underpinning ER under salinity than under PNP stress in our system.

### Stress selects for reversion to metabolic autonomy and mutualism breakdown

Next, we wanted to know if the community composition was altered as a result of the putative genetic changes that may underlie the observed evolutionary rescue. We tested for the presence of each strain at the end of the experiment using allele-specific PCR, looking for the positive amplification of the *ilvA* or *metA* genes, respectively (see Methods). We first confirmed that both strains coexisted until the end of the experiments under stress-free conditions. However, only the ΔI strain survived in all end-point populations subjected to either stress (Figure 3a). The ancestral strains lacked known phages or plasmids that may mediate the horizontal transfer of genes between the two auxotrophs, so the absence of the *ilvA* gene in our populations suggests that evolutionary rescue was achieved by rapid reversion to metabolic autonomy. To confirm this possibility, we cultured these populations in M9 liquid media lacking isoleucine. All populations grew without any supplemented isoleucine (Figure 3b), indicating the acquisition of some compensatory mutations that lead to the reversion from auxotrophic to prototrophic states.

**Figure 3.**
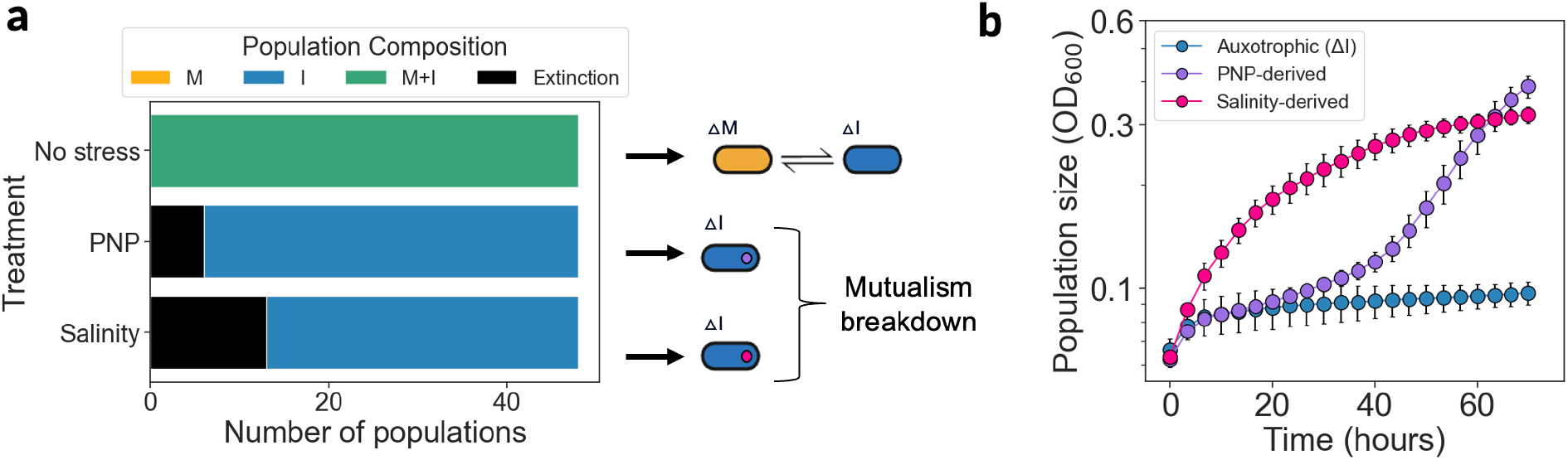
Only a single strain survives the stress and reverts to metabolic autonomy. **a**, Strains present at the end of the experiment. Colors indicate the presence of each strain in the communities as detected by PCR. **b**, Growth curves of recovered populations and of the auxotrophic ancestor of the ΔI strain in M9 without isoleucine addition. Different colors indicate the treatment from which the populations were isolated (purple from PNP, red from salinity and blue is the ancestor ΔI strain). Dots and error bars indicate the mean ± SD of measurements from three technical replicates of each of three evolutionary replicates (i.e. three different populations that were evolved in parallel during the evolutionary rescue experiment). Grow curves of individual replicates are included in supplementary Figure S2.

### Mutualists can survive lower stress levels than the prototrophic strains

Since the prototrophic strain was less affected by either salinity or PNP and did not decline toward extinction during the experimental evolution (Figure 2a), we hypothesized it is less sensitive to stress than the mutualism. To test this hypothesis, we exposed monocultures of the prototroph and cocultures of auxotrophic strains to several stresses with different modes of action: salinity, PNP, hydrogen peroxide, and the antibiotic spectinomycin. The mutualism grew slower than the prototroph under non-stress conditions (Figure S3, mean exponential growth rate of 0.07 ± 0.01 *vs*. 0.04 ± 0.01 (h^-1^); two sided Mann–Whitney U test, p-value ∼ 10^-7^). However, they grew by a similar amount after 48 hours (Figure S3, mean number of doublings 3.1 ± 0.01 *vs*. 2.8 ± 0.6; two sided Mann–Whitney U test, p-value = 0.07). Growth rates and yields declined as stress levels increased, with the mutualism consistently showing lower growth rates and yields than the prototroph across all four types of stress (Figures 4, S4-5). Our results are consistent with previous experiments that reported heightened sensitivity of mutualistic interactions to stress conditions^23,28,29^ and indicate that the auxotrophic strain is able to survive stress levels that would drive the mutualism toward extinction.

**Figure 4.**
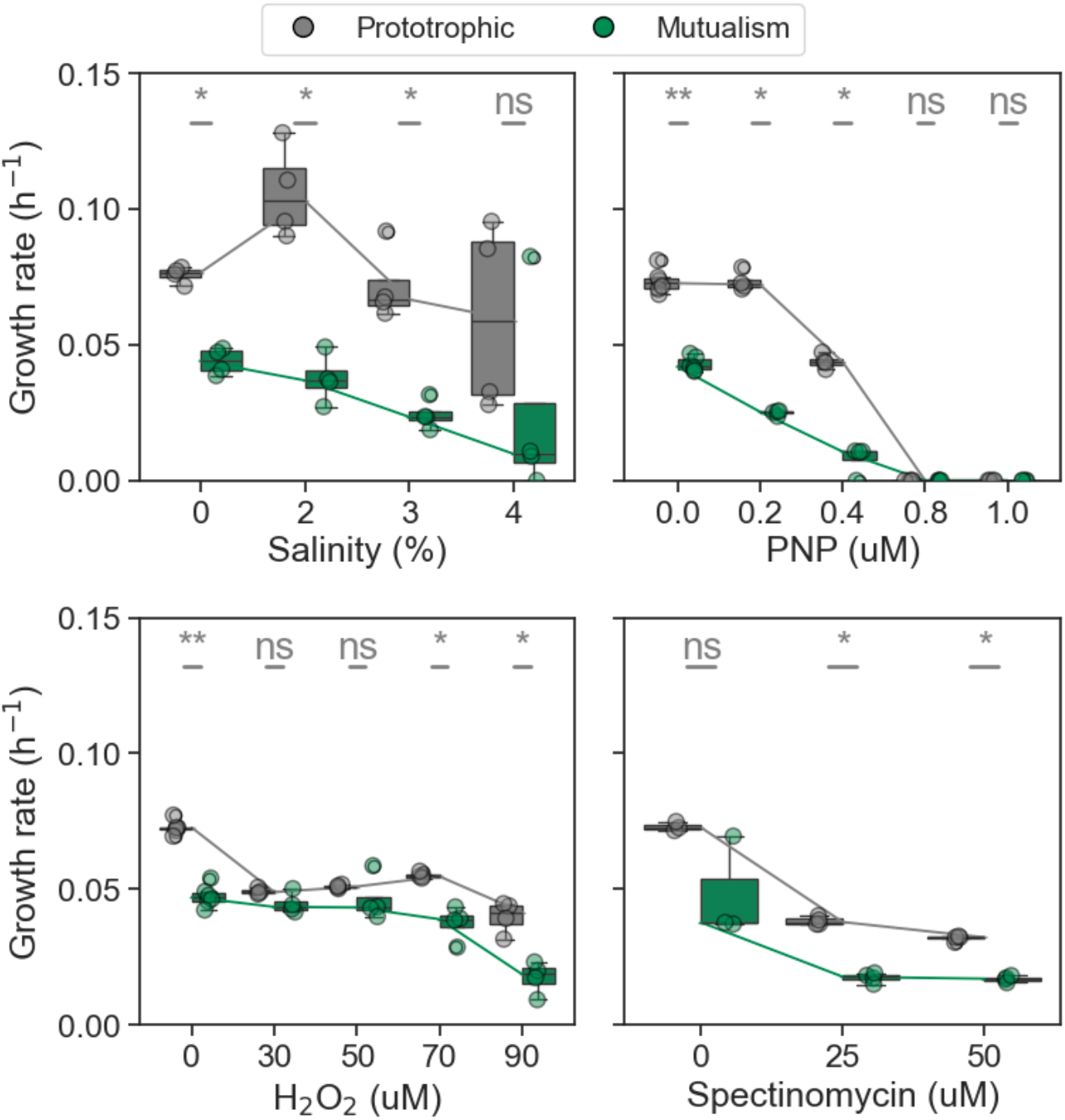
The obligate mutualism is more susceptible to environmental stress than the prototroph. Each panel shows the growth rates of prototrophic (gray) and mutualistic community (green) under different stressors: salinity (%), p-nitrophenol (PNP, µM), hydrogen peroxide (H_2_O_2_, µM), and spectinomycin (µM). Each box plot displays the interquartile range (IQR) of the data, with the horizontal line inside the box indicating the median. The whiskers extend to 1.5 times the IQR, showing the range of the data distribution. There are four technical replicates for salinity, PNP and hydrogen peroxide and three biological replicates for Spectinomycin. The lines connecting the boxplots in each panel indicate the median growth rates for the prototrophic and mutualistic groups across different conditions.

### Mutations in genes involved in amino acid biosynthesis drive the evolutionary rescue

Since the prototrophic populations were less sensitive to either salinity or PNP (Figure 4), we hypothesized that ER in our system was due to bypassing the auxotrophy, rather than stress-specific adaptation. To test this hypothesis, we quantified the growth of strains isolated from the recovered populations when exposed to each of the individual stressors.

We observed that the growth rate of the rescue strains was higher in almost all the cases than that of the mutualism when exposed to either stress (Figure 5a). Furthermore, both the growth rate and the yield of the rescue strains was similar regardless of their evolutionary history when exposed to either condition (Figure S6), indicating a substantial degree of cross-resistance despite the marked differences in the nature of both stresses (osmotic imbalance *vs*. membrane disruption). The fact that the strains recovered from one stress did not outperform those evolved under the alternative stress lends support to our hypothesis that ER was driven mostly by bypassing the auxotrophy.

**Figure 5.**
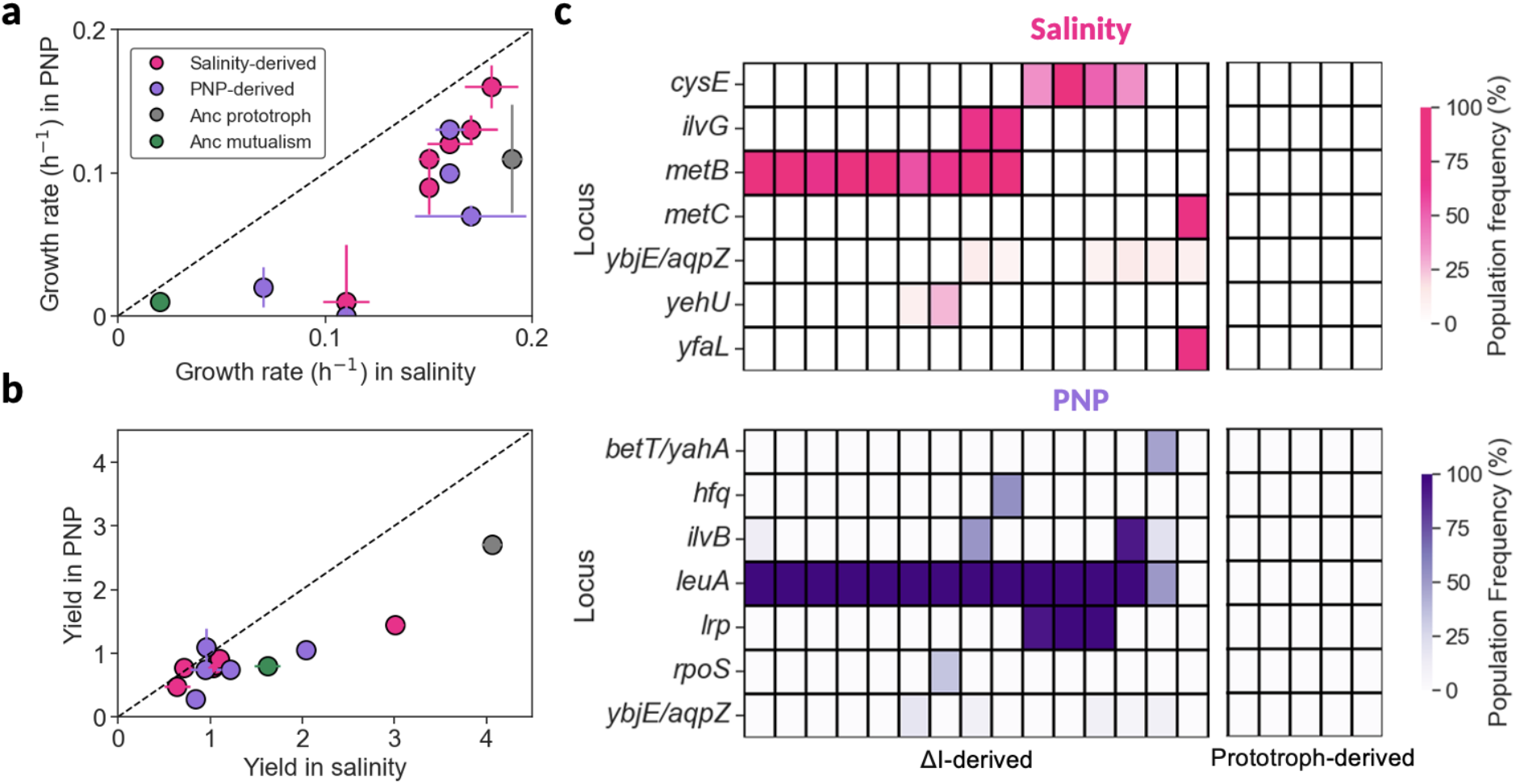
Metabolic pathway mutations drive evolutionary rescue and mutualism breakdown. **a**, Growth rates and **b**, yields of the mutualism, the prototrophic ancestor, and strains evolved in salinity or PNP. For this experiment, we included one population of the prototrophic strain (gray), six evolutionary replicates derived from salinity stress and five from PNP stress, and the ancestral M and I strains that constituted the mutualism. The data are presented as the mean ± SEM (n = 4). (C, D) Genes with mutations present in at least two replicate populations or occurring at a frequency > 0.3, represented by different colors (red for salinity and purple for PNP). Color shades indicate the frequency of each mutation, with darker shades indicating higher mutation frequencies. Specific mutations are detailed in Tables S2-3.

To identify the mutations and putative targets of selection underpinning the observed ER, we subjected the recovered populations to whole-population, whole-genome sequencing. We identified a total of 51 mutational events (22 and 29 under salinity and PNP, respectively) of various types within the derived strain collection (Table S2-3). The most common alterations were non-synonymous single-nucleotide polymorphisms (SNPs), which made up 80% of the total mutations. Specifically, GC→AT and AT→GC transitions were particularly prevalent, accounting for 29% and 31% of all base substitutions, respectively (Figure S7). This aligns well with previous reports of their predominance in bacterial mutation patterns^30^, suggesting that the environmental stresses we used did not introduce significant biases in the overall mutation rates^31,32^.

Moving on to the presumed targets of adaptation, our results show that most high-frequency mutations are related to amino acid metabolism, further supporting the idea that the primary driver of the rescue was the reversion to autonomy rather than stress-specific adaptation. We identified several mutations in metabolic genes that appear to compensate for the loss of *ilvA* gene function in the salinity-evolved populations (Figure 5C, Table S2). In most cases, a putative metabolic mechanism underlying the compensatory phenotype can be identified. The most direct example occurs in the case of *metB*, the most prevalent target of mutation detected. MetB normally produces L-cystathionine and succinate from L-cysteine and O-succinylhomoserine in the methionine biosynthesis pathway. However, when the levels of the substrate L-cystein are low, this enzyme displays a secondary activity, catalyzing the generation of 2-oxobutanoate^33^. This compound is precisely the main product of IlvA, and represents the first step in the synthesis of isoleucine. It is therefore reasonable to speculate that the mutants of MetB we detected could potentially enhance the secondary activity of the enzyme and generate sufficient 2-oxobutanoate so as to bypass the blockade of the first step in isoleucine biosynthesis of the *ΔilvA* background^34^.

A related argument can be proposed for the second most prevalent target of mutation, *cysE*. This enzyme carries out the first step in the pathway of cysteine biosynthesis, so a putative mechanism is that *cysE* mutants result in lower concentration levels of L-cystein, which are the conditions in which wild-type MetB favors the production of 2-oxobutanoate. Along this line, we detected one case that seems explained by mutations in MetC. This enzyme normally catalyzes the reaction step that follows the action of MetB in the methionine biosynthetic pathway. However, this enzyme can also exhibit a secondary activity, converting L-cysteine to 2-aminoacrylate and hydrogen sulfide^35^. It is tempting to hypothesize that the MetC mutant observed favors this reaction, reducing overall L-cystein levels and therefore potentiating the generation of 2-oxobutanoate by MetB. Finally, one further evidence in favor of these explanations is that these three genes are present in 14/15 of the sequenced populations, and most importantly, when mutations are found in one of them, none are found in the others, suggesting that these mutations indeed mediate redundant ways to bypass the isoleucine auxotrophy.

In the PNP treatment, 14 out of 15 sequenced populations had mutations in *leuA* gene (Figure 5c, Table S3), which catalyzes the first step in leucine biosynthesis. The enzyme LeuA is known to have promiscuous activity toward alternative substrates 2-ketobutyrate^36^ and (S)-2-keto-3-methylvalerate^37^, the first and last step in the isoleucine biosynthesis pathway. A tantalizing possibility is that the mutations detected may make the reverse reactions more likely to occur for a given concentrations of substrates and products, as production of either 2-ketobutyrate or (S)-2-keto-3-methylvalerate would suffice to bypass the blockade caused by the *ilvA* knockout^38–41^.

## Discussion

Since we exposed an obligate mutualism to abrupt stress, we expected that evolutionary rescue would be 1) rare and 2) mediated by two different mechanisms: either stress adaptation of both strains, or one strain adapting both to the stress and becoming able to grow independently (Figure 1). We found that evolutionary rescue occurred frequently in our system, likely in large part due to the fact that reverting to metabolic autonomy appears to suffice to also overcome the salinity or PNP stress. Notably, in all the recovered populations it was the same strain, ΔI, that avoided extinction. This may indicate that strain ΔI has a higher mutation rate or larger number of potential mutations that allow it to bypass its auxotrophy compared to the ΔM strain^42^. Alternatively, ΔI may have a larger population than ΔM within the mutualism, resulting in it having a larger overall mutation supply and higher chances to evolve and recover.

Our results suggest that the ability to bypass auxotrophy could be a key factor of evolutionary rescue in obligate mutualism communities, but it is still unclear why bypassing the auxotrophy also led to lower sensitivity to the salinity or PNP stress. It is possible that by bypassing auxotrophy, organisms can conserve vital energy or other cellular resources that would otherwise be expended on acquiring or producing essential nutrients from their mutualistic partners. However, this hypothesis is inconsistent with another experiment in which we found that the salinity sensitivity of the ΔI strain was not relieved by isoleucine supplementation (Figure S7). Moreover, while the rescued populations showed reduced sensitivity to both stresses (Figure 5a,b), the specific mutations that underlie their recovery did depend on the stress in which they evolved. These results suggest a complex interplay between cellular metabolism and stress adaptation, similar to those recently described for the evolution of aminoglycoside resistance^43^.

While evolutionary rescue via stress-induced mutualism breakdown and extinction of one of the partners occurred in our system, different outcomes are likely in other settings. Mutualism breakdown may be less frequent when reverting to autonomy does not result in lower stress sensitivity, as ER would then require two different adaptations. Even when reverting to autonomy leads to stress adaptation, the outcome of stress exposure would depend on the rates with which the mutualistic partners can do so. In our system, a mismatch between the rates at which the strains could bypass their auxotrophies likely contributed to the breakdown of the mutualism. A lower rate of auxotrophy bypass would likely have led to fewer populations recovering. In contrast, if both the ΔI and ΔM strains could readily bypass their auxotrophies, recovery rates would be higher but the outcome more varied - rather than the only the ΔI strain recovering, either one of the strain or both of them could survive.

Further research is needed in order to gain a broader understanding of the factors affecting the likelihood and dynamics of evolutionary rescue of mutualisms. Here, we focused on a mutualism based on amino acid exchange between two strains of the same species. The extent to which our findings extend to other mechanisms of co-dependence, involving more distinct species, and in more natural settings remain unclear. Nonetheless, real-word instances of stress-induced mutualism breakdown have been observed, including breakdown of cooperation in host-microbiota associations exposed to antibiotic treatments^44^, as well as in coral bleaching events^21^. Given the prevalent environmental changes instigated by anthropogenic activities, it is crucial that we gain a deeper understanding of how mutualistic and other ecological interactions affect a community’s adaptive potential and long-term persistence in changing environments.

## Methods

### Strains

*Escherichia coli* used in this study were based on the EcNR1 *E. coli* derivative of MG1655. We used the auxotrophic strains Δ*metA* (hereafter referred to as ‘ΔM’), Δ*ilvA* (hereafter referred to as ‘ΔI’) and the WT (hereafter referred to as prototrophic). The amino acid auxotrophs were generated by Red-recombineering with a chloramphenicol resistance cassette. Information and strains were obtained from Mee et al.^27^. We chose these strains because they have been previously demonstrated to reciprocally exchange amino acids and engage in obligate mutualism.

### Culture conditions

Evolutionary experiments were performed in 96 deep-well plates (maximal volume: 1 ml, Thermo Scientific Nunc) with M9-glucose media [1X M9 salts supplemented with 2 mM MgSO_4_ · 7H_2_O, 0. 1mM CaCl_2_, 1X trace metal solution (Teknova), 0.083 nM thiamine, 0.25 μg/L D-biotin, and 1% (wt/vol) glucose] at 30°C and were shaken at 900 rpm for 48h.

### Population size and growth rate measurements

To determine the growth rate of all populations, growth kinetic experiments were performed with monocultures of the prototrophic strain and cocultures of both auxotrophic strains (ΔI and ΔM). Growth experiments were performed in 96 well plates (200 ul). The optical density was measured in two automated plate readers simultaneously, Epoch2 microplate reader (BioTek) and Synergy microplate reader (BioTek), and was recorded using Gen5 v3.09 software (BioTek). Plates were incubated at 30 °C with a 1 °C gradient to avoid condensation on the lid and were shaken at 250 rpm. OD_600_ was measured every 5 min. Each strain was measured in four technical replicates.

Before calculating the yield and growth rate, a smoothing process was applied to the data. This involved averaging measurements taken every 5 minutes over 8 data points. The yield was computed as the logarithm of the ratio between the final OD_600_ (OD_f_) and the initial OD_600_ (OD_i_) concentration:

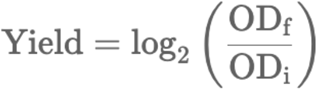

The Growth rate (h^-1^) was calculated as:

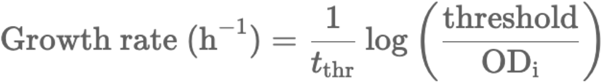

where a threshold (*threshold*) was defined as 1.5 times the initial OD concentration.

### Toxicity assays

To determine the toxicity of several environmental stresses, growth kinetic experiments were performed, as described before, with monocultures of the prototrophic, ΔI and ΔM as well as cocultures of both strains. The assay was performed in in a 96-well plate which was filled with fresh M9 with ot without amino acids and supplemented with increasing amounts of salinity (0, 1, 2, 3, 4% NaCl), PNP: p-nitrophenol (0, 0.2, 0.4, 0.8, 1 mM), hydrogen peroxide (0, 30, 50, 70, 90 H_2_O_2_), Spectinomycin (0, 25, 50 ug/mL) and inoculated with approximately 10^5^ bacteria per well. The plate was incubated for 48 hours at 30°C with shaking.

### Evolutionary rescue experiment

During the evolution experiment, monocultures of the prototrophic and isoleucine- and methionine-auxotrophic strains were grown without any externally supplied amino acid. All the strains involved in the evolution experiment started from one cryogenic stock. Before of the experiment, strains were first picked from an overnight colony into LB–Lennox medium (10 g/L bacto tryptone, 5 g/L NaCl, 5 g/L yeast extract) with chloramphenicol (20 μg/mL, Sigma cat# C0378). After 24h, late–exponential-phase cells were harvested and washed twice in M9 salts (6 g/L Na_2_HPO_4_, 3 g/L KH_2_P_4_, 1 g/L NH_4_Cl, 0.5 g/L NaCl). Then, all cell concentrations were adjusted to 0.1 OD_600_ using M9 media. Coculture growth was performed by equal-volume inoculation of each strain at a seeding OD_600_ of 0.05. Precultures were then used to inoculate 46 replicates for each of the two cultures: (1) monoculture of prototrophic and (2) coculture of ΔI and ΔM. Cultures were grown in a volume of 800 µl in 96-well plates at 30°C and were shaken at 900 rpm for 48h. 40 µl of the resulting cultures were transferred every 48 into 760 µl of fresh medium. During the first three growth cycles, no stress treatment was applied to the cultures to allow populations to equilibrate. At the fourth transfer, each test culture was split up into three different environments: control (no stress), salinity (3% NaCl) and PNP (0.4 mM). Growth of all cultures was tracked by quantifying their population density (OD_600_), while propagating them to fresh medium with the specific stress.

These two stresses were chosen to maximize differences in terms of mode of action, and are relevant pollutants in the environment. Salinity exerts its influence primarily through osmotic stress, where high salt concentrations challenge microbial cells by disrupting their osmotic balance. On the other hand, PNP acts as a direct chemical stressor by interfering with critical cellular processes, including enzyme functions.

### Dynamics of the U-shaped ER curve

Before performing the any calculation, we smooth the data by applying a moving average smoothing technique with a window size of three transfers.

The *Rescue Transfer* is defined as the transfer when an experimental culture exhibits the first increase in population size (OD_600_) after being exposed to environmental change, either salinity or the presence of PNP.

The *Maximum recovery rate* is defined as the maximal positive change in population size (OD_600_) per transfer for each experimental culture after the exposure to stress.

### Sequencing and genomic analyses of rescue populations

Each ∼20 generations all communities were frozen at ™80 °C with 50% glycerol in a 96-deep well plate. In order to do population sequence, we use a loop and resuspend in 3 mL LB at 30°C, overnight with shaking. We extracted genomic DNA using the Norgen Biotek Corp Bacterial Genomic DNA Isolation Kit (Cat No. 17900) and sent it to the sequencing facility at SEQCENTER (PA, USA, https://www.seqcenter.com/). The samples were sequenced on an Illumina NextSeq 2000. Demultiplexing, quality control, and adapter trimming were performed by the sequencing center with bcl-convert. Average coverage for short reads is 121 with a standard deviation of 40. Analysis of the NGS sequencing data from samples was performed using breseq (v.0.37) software tool^45,46^. This tool is free-access, based on Linux, and has been extensively used to analyze genome’s sequencing data. To prepare the input for breseq, we first extract fastq.gz files containing raw sample genomic data into fastq files. These files were then used for the analysis. In addition, we used the *Escherichia coli* str. K-12 substr. MG1655, complete genome genebank (.gbk) extension file as the reference sequence, since it has already been annotated. Additionally, we remove possible mutations due to our ancestor by comparing the complete genome sequence with the sequence of our ancestor. We run breseq in a population default mode without any modifications.

## Acknowledges

We thank Valeria Tzvichenko and Rotem Shner for the laboratory assistance, and members of the Friedman lab and Couce lab for helpful discussions. JF was supported by the Israel Science Foundation (grant No. 883/22). AC supported by the Agencia Estatal de Investigación (Centros de Excelencia “Severo Ochoa”, SEV-2016-0672 and CEX2020-000999-S), and a Comunidad de Madrid “Talento” Fellowship (2019-T1/BIO-12882). IJMJ was supported by “Margarita Salas” post-doctoral fellowship (Universidad de Málaga, Unión Europea–NextGeneration EU, Ministerio de Universidades, Spain).

## Author contributions

I.J.M.J, A.C. and J.F. designed the study. I.J.M.J., Y.S. and A.M. performed the experiments. I.J.M.J, A.C. and J.F. performed the analysis. I.J.M.J, A.C. and J.F. wrote the manuscript.

