## Supplementary material for "Reversion to metabolic autonomy underpins evolutionary rescue of a bacterial obligate mutualism": SI

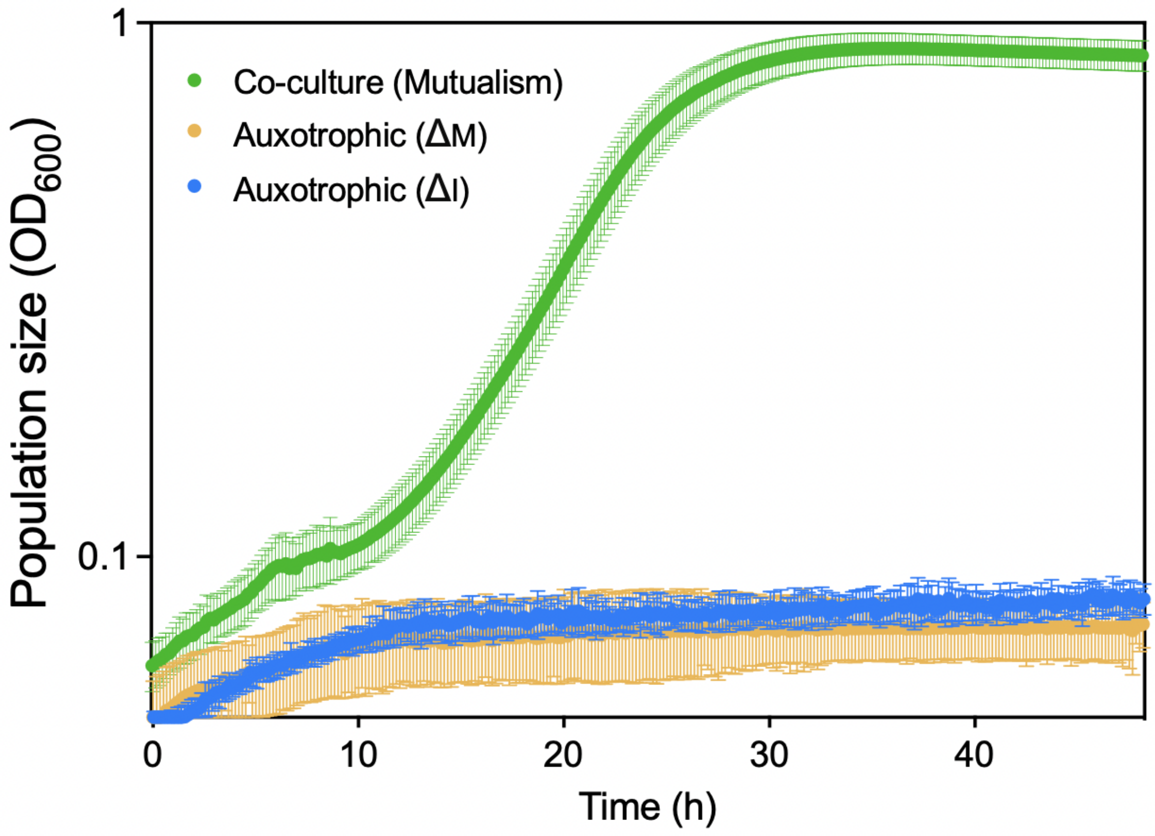


**Figure S1. Obligate metabolic cross feeding of auxotrophic strains.** The growth curves depict the E. coli auxotrophic strains ΔM, ΔI, and their coculture. The data are presented as the mean ± SD (n = 8).

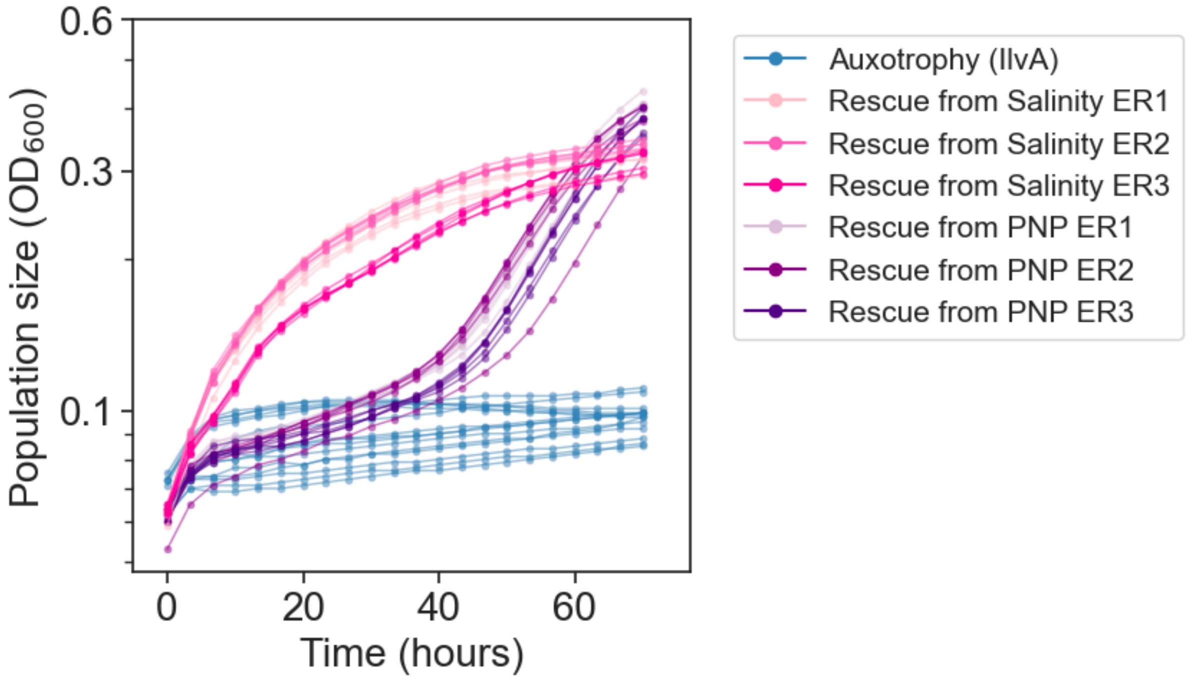


**Figure S2. Rescue strains can grow without isoleucine supplied.** Growth curves of recovered populations and of the auxotrophic ancestor of the ΔI strain in M9 without isoleucine addition. Different range of colors indicate the treatment from which the populations were isolated (purple from PNP, pink from salinity, and blue is the ancestor ΔI strain). ER denotes a evolutionary replicate.

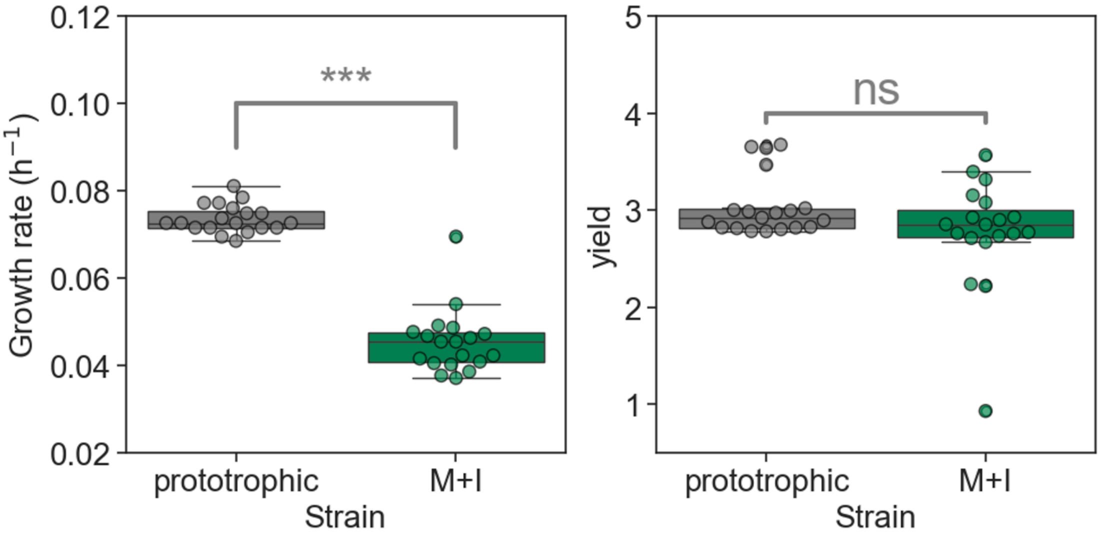


**Figure S3. Comparison of Growth Rate (h$^{-1}$) and yield between prototrophic and ΔM+ΔI (mutualism) Strains without stress.** Boxplots shows growth rate and yield of the prototrophic and ΔM+ΔI strains. Each box plot displays the interquartile range (IQR) of the data, with the horizontal line inside the box indicating the median. The whiskers extend to 1.5 times the IQR, showing the range of the data distribution.

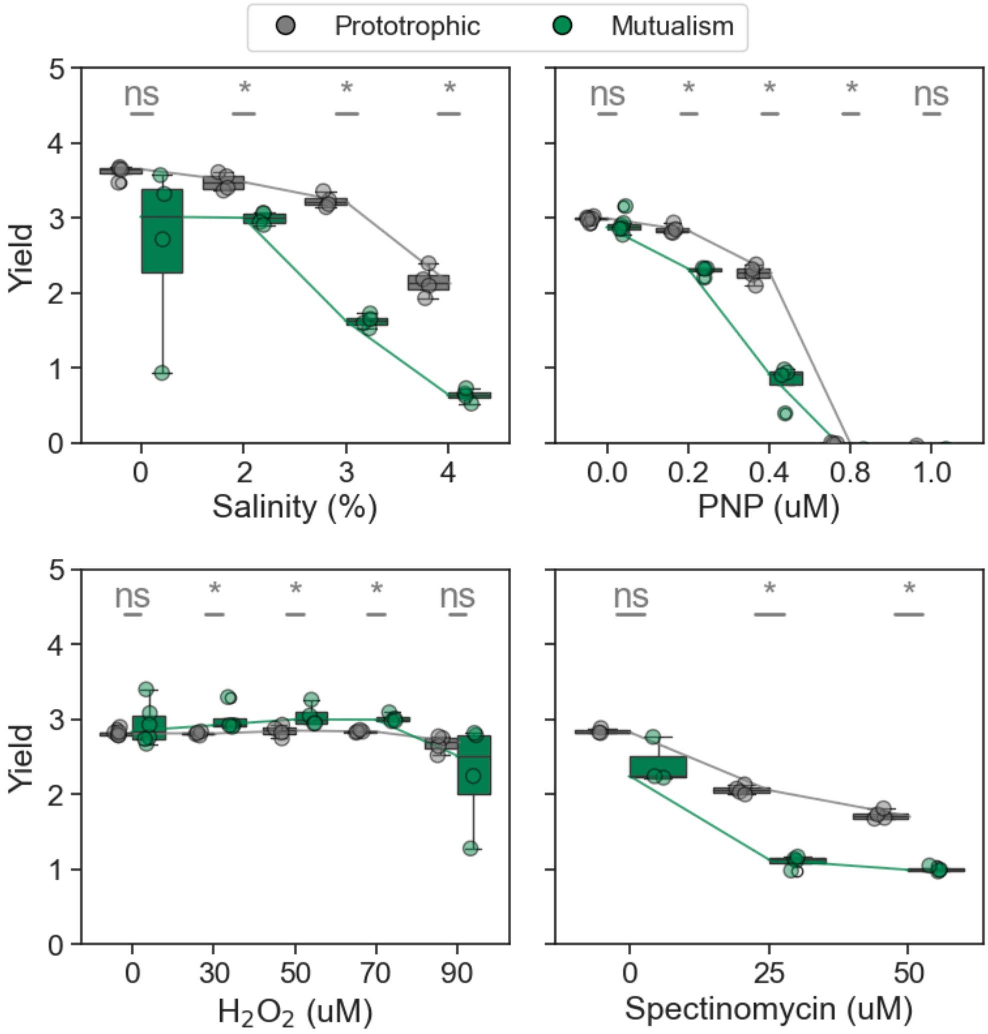


**Figure S4. The obligate mutualism is more susceptible to environmental stress than the prototroph.** Each panel shows the yield of prototrophic (gray) and mutualistic community (green) under different stressors: salinity (%), p-nitrophenol (PNP, µM), hydrogen peroxide ($H_2O_2$, µM), and spectinomycin (µM). Each box plot displays the interquartile range (IQR) of the data, with the horizontal line inside the box indicating the median. The whiskers extend to 1.5 times the IQR, showing the range of the data distribution. There are four technical replicates for salinity, PNP and hydrogen peroxide and three technical replicates for Spectinomycin. The lines connecting the boxplots in each panel indicate the median yields for the prototrophic and mutualistic groups across different conditions.

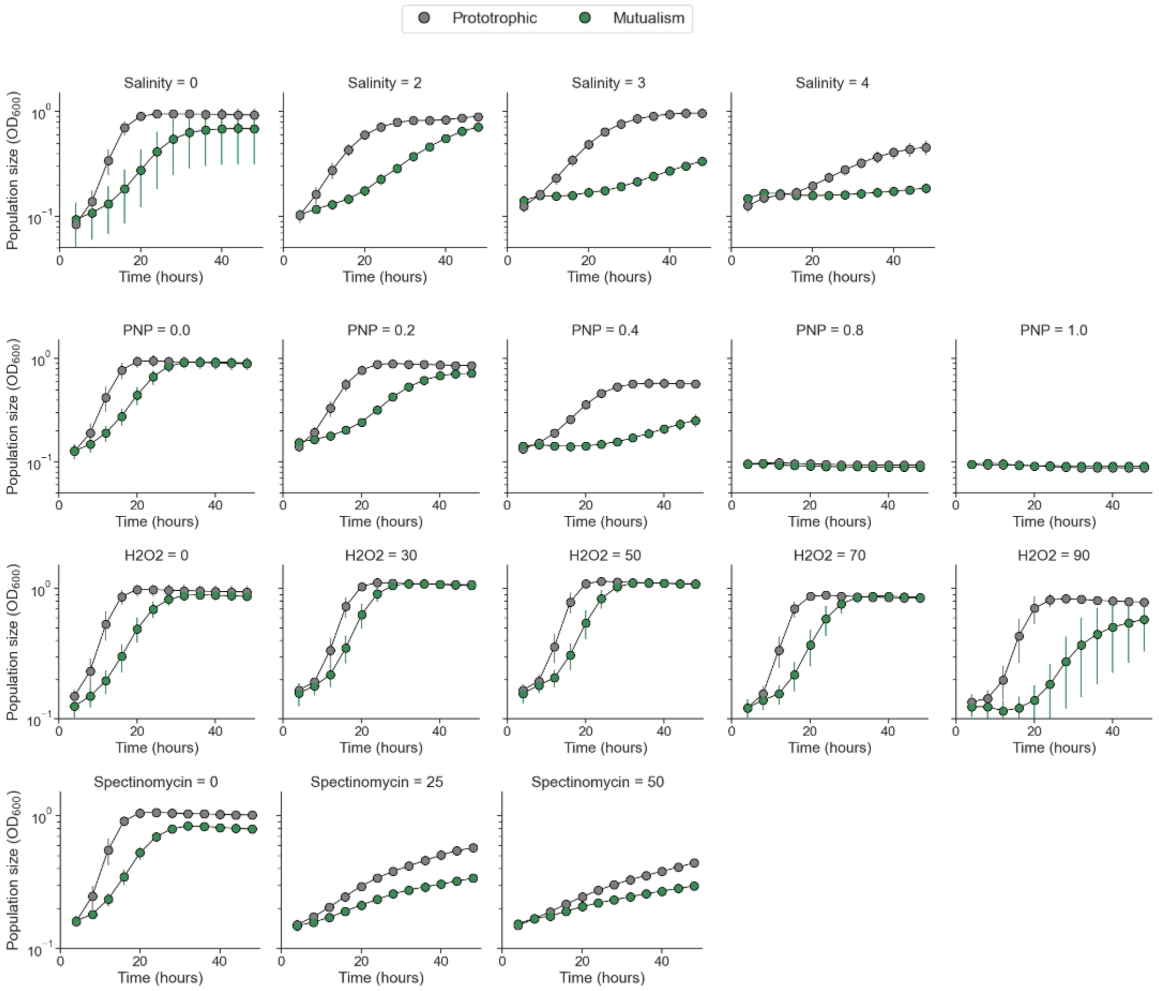


**Figure S5. Mutualism are more sensitive to stress than the prototrophic strain.** Growth curves of the prototrophic strain and the mutualism under non-stress and different stressors: salinity (%), p-nitrophenol (PNP, µM), hydrogen peroxide ($H_2O_2$, µM), and spectinomycin (µM). Data shown the mean ± SD (n=4 for salinity, PNP and H2O2 and n=3 for Spectinomycin).

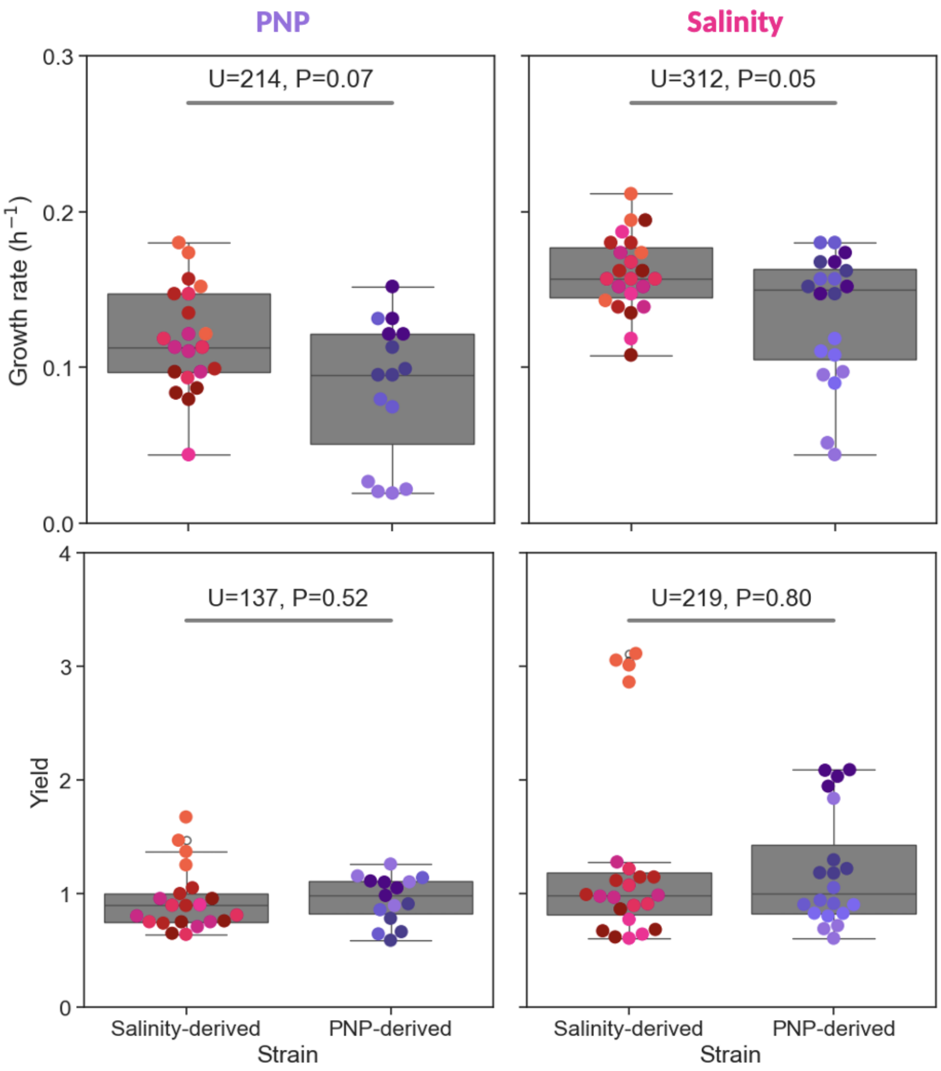


**Figure S6. Growth rate and the yield of the rescue strains are similar regardless of their evolutionary history.** Each column represents the stress condition under which the isolate-derived strains were grown. Each box plot displays the interquartile range (IQR) of the data, with the horizontal line inside the box indicating the median. The whiskers extend to 1.5 times the IQR, showing the range of the data distribution. Points represent different biological replicates (6 for salinity and 5 for PNP) and technical replicates (3) are indicated with different tones of purple (PNP-derived) or pink (Salinity-derived). The statistical test performed was a two-sided Mann–Whitney U test.

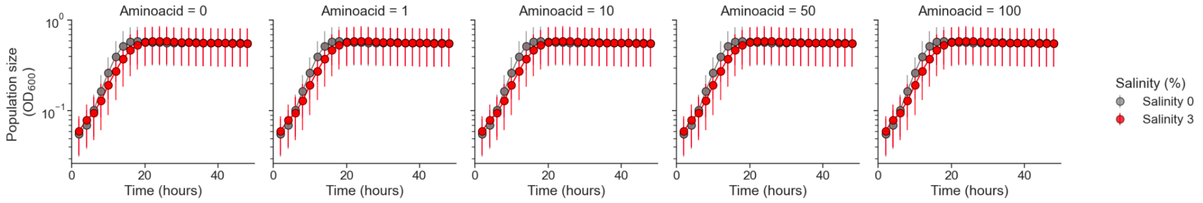


**Figure S7. Sensitivity to salinity is not affected by amino acids content for the prototrophic strain.** Growth curves of the prototrophic strain without salinity and with 3% salinity at different levels of isoleucine supplemented to the media. Each panel shows the growth curve at different isoleucine concentration, as indicated by the panel title (gray and red circles indicate 0% 3% salinity, correspondingly). The data are presented as the mean ± SD (n = 4)

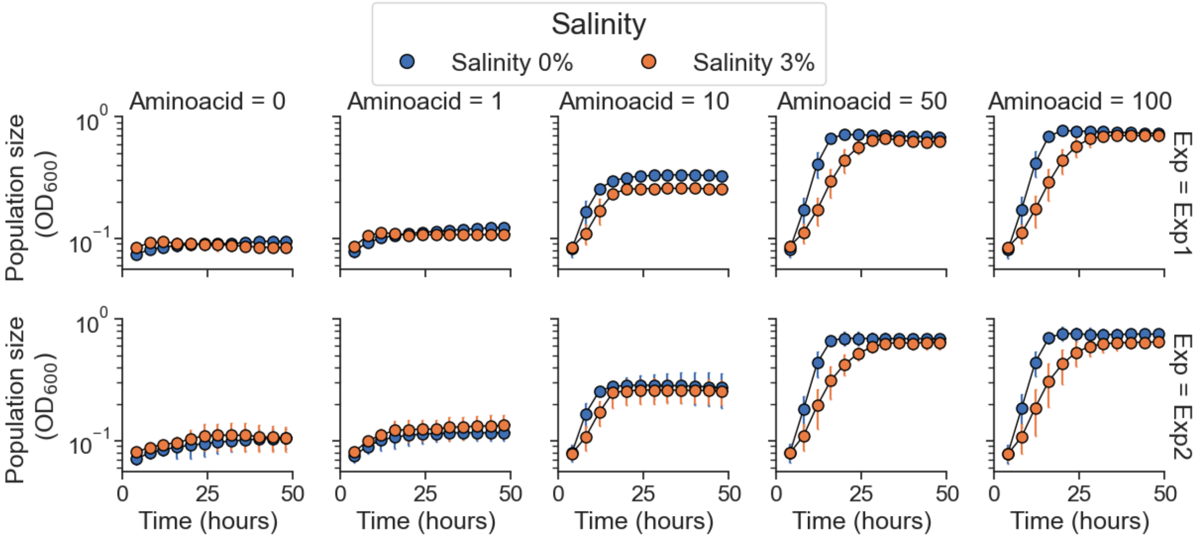


**Figure S8. Sensitivity to salinity is not affected by amino acids content for the auxotrophic ΔI strain.** Growth curves of the auxotrophic ΔI strain without salinity under non-stress and with 3% salinity (3%) at different levels of isoleucine supplemented to the media. Each row corresponds to one of two independent experiments. Each panel depicts the growth curve at different isoleucine concentration (blue circles for 0% and orange circles for 3% salinity). The data are presented as the mean ± SD (n = 4)

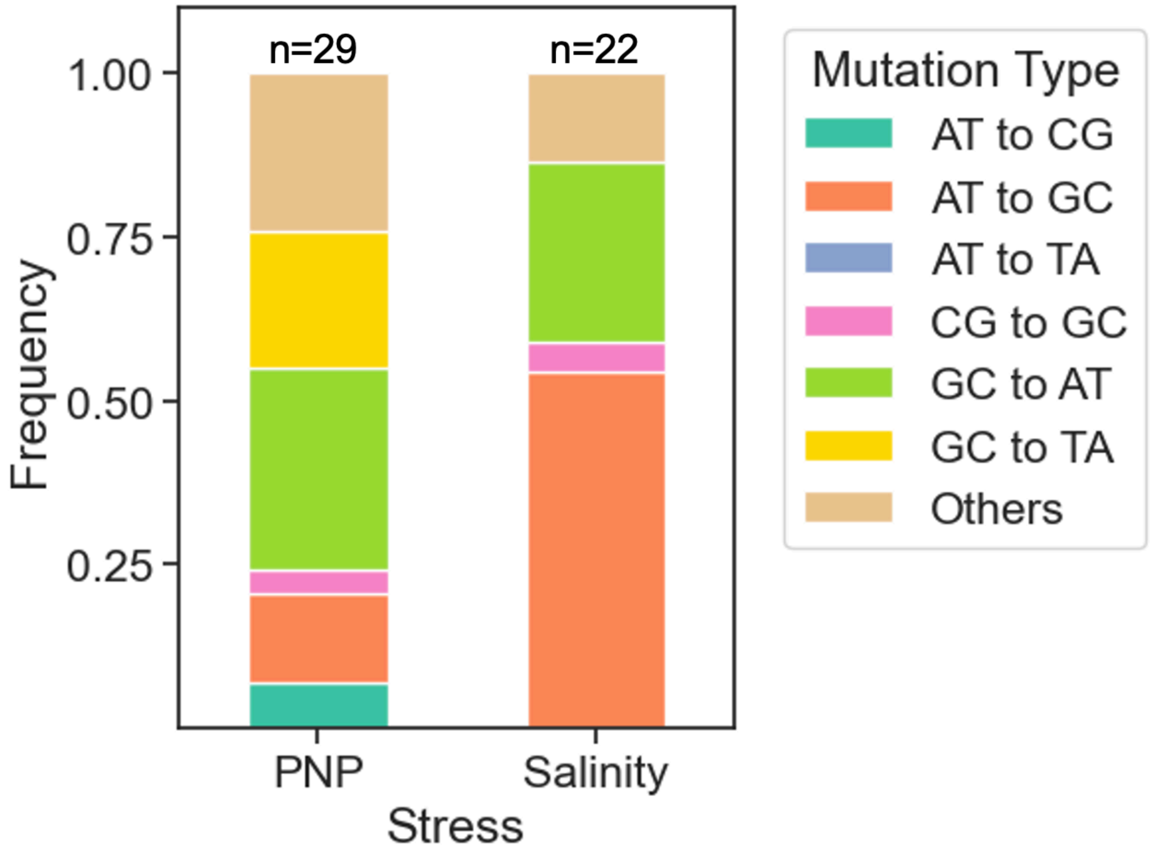


**Figure S9. Environmental stresses we used did not introduce significant biases in the overall mutation rates.** Base substitution frequencies observed in the genomes of strains evolved under salinity or PNP stress. Colors indicate the six possible substitutions. The specific mutations are listed in Tables S1-2.

**Table S1.** List of primers used in this study

| Gene | Primer sequence |
| --- | --- |
| ilvA_vf | GTGAACGTCAGGTCTCCTTTG |
| ilvA_vr | TATGCGCCGCGCAGC |
| metA_vf | TCACCTTCAACATGCAGGC |
| metA_vr | ATATCGACCTGCAAAGGTGAGT |

**Table S2.** List of mutations found in the ΔI strain derived from the evolutionary rescue experiment with salinity.

| Locus | mutation | annotation | population | description |
|---|---|---|---|---|
| *cysE* ← | C→T | A28T (GCC→ACC) | 3 | serine acetyltransferase |
| | G→A | R125C (CGT→TGT) | 5 | |
| | A→G | Y31H (TAC→CAC) | 7 | |
| | IS*1* (+) +9 bp | coding (780-788/822 nt) | 11 | |
| *ilvG* → | (AAATC)$_{1\rightarrow2}$ | pseudogene (979/984 nt) | 13 | pseudogene, acetohydroxy acid synthase II; acetolactate synthase II, large subunit, cryptic, interrupted |
| | Δ1 bp | \|pseudogene (1/663 nt) | 14 | |
| *metB* | A→G | N15S (AAT→AGT) | 2 | cystathionine gamma-synthase, PLP-dependent |
| | A→G | N15S (AAT→AGT) | 6 | |
| | C→T | T322M (ACG→ATG) | 8 | |
| | C→T | T322M (ACG→ATG) | 9 | |
| | (TTACTC)$_{1\rightarrow2}$ | coding (140/1161 nt) | 10 | |
| | A→G | N15S (AAT→AGT) | 12 | |
| | A→G | N15S (AAT→AGT) | 13 | |
| | A→G | N15S (AAT→AGT) | 14 | |
| | A→G | N15S (AAT→AGT) | 15 | |
| *metC* → | A→C | D185A (GAC→GCC) | 4 | cystathionine beta-lyase, PLP-dependent |
| *ybjE* ← / ← *aqpZ* | T→C | intergenic (-369/+126) | 1 | putative transporter/aquaporin Z |
| | C→G | intergenic (-341/+154) | 7 | |
| | T→C | intergenic (-369/+126) | 11 | |
| | T→C | intergenic (-369/+126) | 13 | |
| | T→C | intergenic (-369/+126) | 14 | |
| *yehU* | G→A | L375F (CTT→TTT) | 8 | putative sensory kinase in two-component system with YehT, inner membrane protein |
| | G→A | T325I (ACC→ATC) | 10 | |
| *yfaL* ← | Δ18 bp | coding (2791-2808/3753 nt) | 4 | adhesin |

**Table S3.** List of mutations found in the ΔI strain derived from the evolutionary rescue experiment with PNP.

| Locus | mutation | annotation | population | description |
|---|---|---|---|---|
| *betT*→/→*yahA* | T→C | intergenic (+386/-489) | 12 | 2-isopropylmalate synthase |
| *hfq*→ | G→T | K31N (AAG→AAT) | 5 | global sRNA chaperone; HF-I, host factor for RNA phage Q beta replication |
| *ilvB*← | +A | coding (623/1689 nt) | 6 | acetolactate synthase I, large subunit |
| | G→T | S8* (TCG→TAG) | 14 | |
| *leuA*← | C→T | A460T (GCG→ACG) | 1 | 2-isopropylmalate synthase |
| | G→A | R268C (CGC→TGC) | 3 | |
| | G→A | A453V (GCC→GTC) | 4 | |
| | C→T | A460T (GCG→ACG) | 5 | |
| | C→T | G427D (GGT→GAT) | 6 | |
| | A→G | S400P (TCT→CCT) | 7 | |
| | G→T | R268S (CGC→AGC | 8 | |
| | C→T | A460T (GCG→ACG) | 9 | |
| | C→A | G462V (GGT→GTT) | 10 | |
| | A→C | S488A (TCT→GCT) | 11 | |
| | Δ12 bp | coding (1356-1367/1572 nt | 12 | |
| | C→A | G462V (GGT→GTT) | 13 | |
| | C→T | A460T (GCG→ACG) | 14 | |
| | A→C | L461R (CTG→CGG) | 15 | |
| *lrp*→ | C→T | P87L (CCG→CTG) | 10 | DNA-binding transcriptional dual regulator, leucine-binding |
| | G→T | R137L (CGT→CTT) | 11 | |
| | C→T | P87L (CCG→CTG) | | |
| *rpoS*← | Δ1 bp | coding (10/993 nt) | 1 | RNA polymerase, sigma S (sigma 38) factor |
| *ybjE*←/←*aqpZ* | T→C | intergenic (-369/+126) | 3 | putative transporter/aquaporin Z |
| | T→C | intergenic (-369/+126) | 12 | |
| | C→G | intergenic (-341/+154) | 10 | |
